# Integrated heart rate variability and physiological profiling reveals autonomic phenotypes in older adults from a high-southern-latitude population

**DOI:** 10.64898/2026.03.27.714667

**Authors:** David Medina-Ortiz, Matias Castillo-Aguilar, Diego Mabe-Castro, Matías Mabe-Castro, Cristián Nuñez

## Abstract

Heart rate variability (HRV) is widely used to assess autonomic regulation, but its interpretation in older adults is influenced by age, sex, body composition, and hemodynamic status, particularly in underrepresented populations living in geographically extreme environments. We analyzed 530 community-dwelling older adults from the Magallanes region in southern Chile using an integrated framework that combined HRV indices with demographic, anthropometric, and cardiovascular descriptors. After quality-controlled preprocessing, we characterized the distribution and association structure of autonomic and physiological variables and then performed a large-scale unsupervised clustering benchmark across multiple feature spaces, dimensionality-reduction strategies, and clustering algorithms. Conventional descriptors explained only a limited proportion of HRV variability, whereas integrated multivariate analysis revealed a structured continuum of autonomic heterogeneity. A six-cluster solution provided the best compromise between separation, balance, and physiological interpretability, identifying profiles that differed in HRV magnitude, blood pressure burden, body composition, sex distribution, and age structure. These findings indicate that autonomic regulation in older adults cannot be adequately summarized by isolated descriptors such as age, body mass index, or blood pressure alone. Instead, it is better represented as a multidimensional physiological organization that supports future hypothesis generation for risk stratification and longitudinal monitoring in aging populations.

## 1. Introduction

Heart rate variability (HRV) is widely recognized as a non-invasive biomarker of autonomic nervous system regulation and cardiovascular adaptability (Agorastos et al., 2023). By quantifying beat-to-beat fluctuations in cardiac cycle duration, HRV provides indirect information about the dynamic interplay between sympathetic and parasympathetic influences on the heart, as well as the organism’s capacity to respond to internal and external demands (Arakaki et al., 2023; Stephenson et al., 2021). Because of these properties, HRV has been extensively used in clinical and physiological research as an indicator of autonomic function, cardiovascular regulation, stress-related responses, and broader physiological resilience (An et al., 2020; Perna et al., 2020; Hourani et al., 2020; Young and Benton, 2018).

A variety of HRV indices have been developed to characterize complementary dimensions of autonomic modulation. Among the most commonly used time-domain metrics are the standard deviation of normal-to-normal intervals (SDNN), which reflects overall variability in cardiac autonomic input (Van den Berg et al., 2018), and the root mean square of successive differences (RMSSD), typically interpreted as a marker of short-term vagally mediated modulation (Minarini, 2020). In the frequency domain, low-frequency (LF), high-frequency (HF), and very-low-frequency (VLF) components are commonly used to describe oscillatory patterns of cardiovascular regulation and autonomic control (Garbilis and Mednieks, 2024).

Although these indices provide valuable information about autonomic function, their physiological interpretation is not always straightforward. HRV measures are influenced by multiple demographic, anthropometric, and cardiovascular factors, including age, sex, body composition, and hemodynamic status. Consequently, the interpretation of HRV values requires considering additional physiological descriptors rather than evaluating HRV indices in isolation, particularly when studying heterogeneous human populations.

This issue is especially relevant in older adults. Aging is associated with structural and functional changes in both the cardiovascular and autonomic nervous systems, including reduced baroreflex sensitivity, altered cardiac responsiveness, vascular stiffening, and shifts in sympathovagal balance (Monahan, 2007). In addition, anthropometric characteristics and blood pressure parameters may substantially modulate autonomic regulation in later life, further increasing inter-individual variability. As a result, HRV values in older populations must be interpreted within the context of demographic, anthropometric, and cardiovascular characteristics in order to obtain a physiologically meaningful understanding of autonomic function (Olivieri et al., 2024).

Despite the extensive use of HRV in both clinical and research settings, many studies continue to examine a limited number of HRV variables or analyze them independently from other relevant physiological measures. This fragmented approach may reduce the biological interpretability of HRV findings, particularly in aging populations, in whom autonomic regulation is shaped by multiple interacting factors. Variables such as age, sex, body mass index, systolic and diastolic blood pressure, pulse pressure, and mean arterial pressure may all influence the autonomic profile reflected by HRV. Therefore, an integrative analytical framework is needed to better understand how HRV metrics relate to the physiological characteristics of the individuals being studied.

Another limitation in the current literature is the relatively limited use of multivariate exploratory approaches for HRV characterization. While group comparisons and isolated associations remain common, fewer studies examine the joint structure of HRV, anthropometric, and cardiovascular variables through integrated descriptive, correlational, and multivariate analyses. Because autonomic regulation is inherently multidimensional, patterns of association across variables may reveal physiologically meaningful profiles that are not evident from univariate analyses alone. A more comprehensive characterization of these relationships may improve interpretation, support the identification of coherent autonomic phenotypes, and provide a stronger basis for future predictive or stratification models (Silva et al., 2018).

Despite the growing use of HRV in aging research, studies often rely on isolated indices or predefined clinical categories, which may obscure latent physiological organization across individuals. This limitation is particularly relevant in older adults, in whom autonomic regulation emerges from the interaction of cardiovascular load, body composition, aging-related functional decline, and sex-related biological differences. In geographically extreme populations, such as those living in southern high-latitude regions, these interacting physiological domains may be further shaped by environmental and behavioral conditions, yet integrated autonomic phenotyping remains scarce. Before advancing toward inferential, predictive, or machine learning approaches, it is necessary to understand the distribution of the variables, their internal consistency, and the structure of their associations within the study population (Mabe-Castro et al., 2024; Cipriani et al., 2026). This is particularly relevant when working with physiological datasets, where data quality issues and inter-individual variability may substantially affect interpretation.

We therefore aimed to determine whether integrated physiological profiling could identify autonomic heterogeneity in older adults from southern Chile beyond conventional clinical classifications. We hypothesized that HRV variability in this population would be only partially explained by isolated descriptors such as age, sex, BMI, or blood pressure, and that multivariate unsupervised analysis would reveal physiologically coherent candidate profiles reflecting latent autonomic organization.

## 2 Results and discussion

### 2.1 Dataset construction and preprocessing quality assessment

The preprocessing pipeline was designed to transform the original spreadsheet into an analysis-ready dataset while preserving traceability of all cleaning and validation steps. The raw file contained 530 observations together with demographic, anthropometric, cardiovascular, and heart rate variability variables obtained during the experimental protocol. Preprocessing included variable harmonization, correction of formatting inconsistencies, verification of numeric values, and generation of diagnostic indicators used to evaluate data quality before statistical analysis. After preprocessing, the dataset retained the complete cohort and contained the full set of variables required for subsequent descriptive and multivariate analyses.

Diagnostic summaries obtained before and after preprocessing indicated that the dataset showed only limited missingness in the main physiological variables. The highest proportion of missing values was observed in HRV metrics, with 11 missing observations for SDNN (2.08%), 8 for RMSSD (1.51%), and 1 for mean heart rate (0.19%), while all remaining variables were complete or nearly complete. No duplicated rows and no fully empty rows were detected, confirming the structural integrity of the dataset. Because missing values affected only a small fraction of the cohort, no participants were excluded during preprocessing.

Internal consistency evaluation revealed a small number of discrepancies in anthropometric and cardiovascular variables. Height values expressed in different units were inconsistent in 9 participants (1.70%), body mass index differed from the value recomputed from cleaned height and weight in 8 participants (1.51%), and mean heart rate calculated from the RR interval did not match the recorded value in 15 participants (2.83%). These discrepancies involved only a small proportion of the cohort and did not justify subject exclusion. All cases were retained and marked with quality-control indicators in order to preserve the original sample while allowing assessment of their potential impact on subsequent analyses.

Range validation showed that only a small fraction of the dataset contained values outside the predefined physiological limits. Fourteen participants (2.64%) presented at least one out-of-range measurement, most of which corresponded to HRV variables, particularly mean RR interval, SDNN, and mean heart rate. In contrast, demographic, anthropometric, and blood pressure variables were almost entirely within the expected ranges. These observations were retained and flagged in the dataset so that their potential influence on the statistical and clustering analyses could be evaluated without reducing the cohort size.

The final curated dataset included all 530 participants with harmonized demographic, anthropometric, cardiovascular, and HRV variables together with preprocessing quality indicators. Only a small proportion of observations showed inconsistencies or out-of-range values, indicating good overall data quality. This dataset was used as the input for the statistical characterization and the unsupervised clustering analyses described in the following sections.

To assess the potential influence of minor preprocessing inconsistencies, sensitivity analyses were performed after excluding flagged observations. The resulting partitions showed no substantive change in the overall multivariable structure of the cohort, suggesting that the main clustering patterns were not solely driven by preprocessing anomalies.

### 2.2 Statistical characterization of HRV and physiological descriptors

A comprehensive statistical characterization of the cohort was performed to describe the distribution of heart rate variability (HRV) metrics together with demographic, anthropometric, and cardiovascular descriptors, and to examine how these variables interact across different physiological domains. The results are summarized in Figure 1, which integrates cohort composition, distributional properties, correlation structure, and nonparametric effect size analyses across single and combined physiological factors, providing a global view of the variability present in the population.

**Figure 1:**
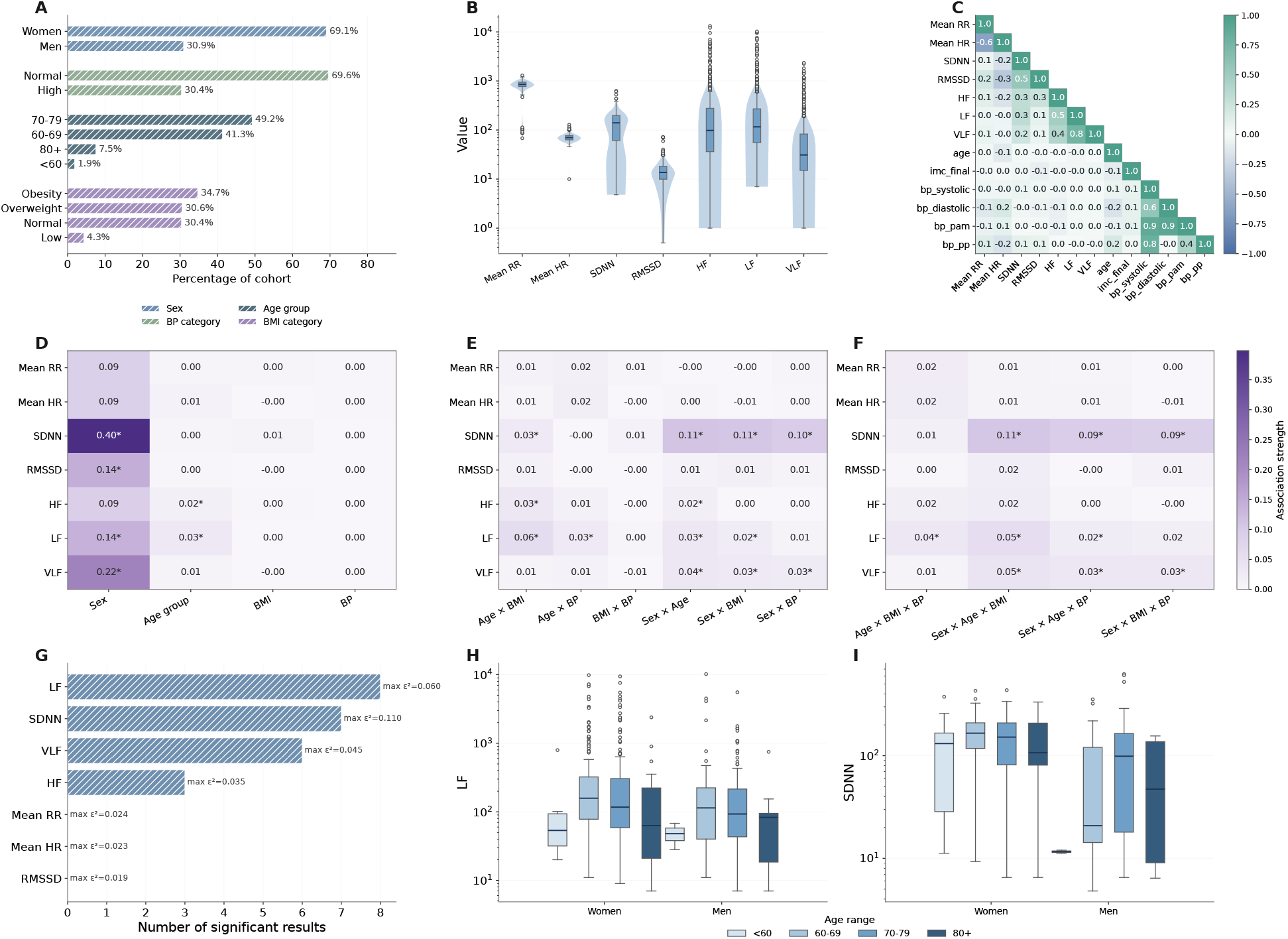
Statistical characterization of HRV metrics and physiological descriptors in the study cohort. (A) Cohort composition across sex, blood pressure category, age range, and BMI category. (B) Distribution of HRV variables (Mean RR, Mean HR, SDNN, RMSSD, HF, LF, VLF) shown using violin and box plots in logarithmic scale. (C) Correlation matrix between HRV metrics and physiological descriptors, including age, BMI, and blood pressure variables. (D) Effect sizes for single-category comparisons (sex, age group, BMI, blood pressure) computed using Cliff’s delta or ε^2^ depending on the number of groups. (E) Effect sizes for double-category comparisons. (F) Effect sizes for triple-category comparisons. (G) Global ranking of HRV variables according to number of significant comparisons and maximum effect size. (H) Distribution of LF across sex and age ranges. (I) Distribution of SDNN across sex and age ranges. Asterisks indicate statistically significant comparisons after multiple-testing correction.

The study population was composed predominantly of women (69.1%) compared with men (30.9%), and most participants were classified in elevated blood pressure categories, with 69.6% categorized as high and 30.4% as normal. The age distribution was centered on older adults, with the largest group corresponding to individuals between 70–79 years (49.2%), followed by 60–69 years (41.3%), while participants aged 80 years or older represented 7.5% of the cohort and individuals younger than 60 years accounted for only 1.9% (Figure 1 A). Regarding body mass index (BMI), most individuals were classified as overweight (30.6%) or obese (34.7%), whereas 30.4% were within the normal range and only 4.3% were underweight. Overall, the cohort is characterized by a high prevalence of cardiometabolic risk factors, a condition expected in older populations and known to influence autonomic regulation, although not necessarily in a uniform manner across individuals.

The distribution of HRV metrics showed marked heterogeneity across variables, with several indices spanning multiple orders of magnitude (Figure 1 B). Mean RR and mean HR presented relatively narrow distributions compared with spectral components, while SDNN, HF, LF, and VLF exhibited wide dispersion and strong right-skewed behavior, consistent with the known variability of frequency-domain HRV measures in aging populations. This heterogeneity indicates that autonomic regulation in the cohort cannot be represented by a single scale of variability and justifies both logarithmic visualization and the use of nonparametric statistical approaches in subsequent analyses. Importantly, the broad dispersion observed in long-term and spectral components suggests the presence of substantial inter-individual differences that are not captured by conventional clinical categories alone.

Correlation analysis revealed a structured but non-redundant relationship between HRV indices and physiological descriptors (Figure 1 C). As expected, mean RR and mean HR were strongly negatively correlated (*r* ≈ − 0.6), reflecting their mathematical dependence. Among HRV metrics, moderate to strong positive correlations were observed between SDNN, HF, LF, and VLF, particularly between LF and VLF (*r >* 0.7), indicating shared contributions of autonomic modulation and overall variability. In contrast, associations between HRV variables and anthropometric descriptors were weak, with correlations close to zero for BMI and height, suggesting that body composition alone does not strongly determine basal autonomic variability in this cohort. Blood pressure variables showed moderate internal correlations, especially between systolic pressure, mean arterial pressure, and pulse pressure, but their association with HRV indices remained limited, indicating that resting cardiac autonomic regulation and hemodynamic state represent partially independent physiological domains. This partial independence supports the idea that multiple regulatory mechanisms contribute to the observed variability and that their combined effects cannot be captured by simple linear relationships.

To quantify the influence of individual physiological factors on HRV, nonparametric effect sizes were computed for single-category comparisons using Cliff’s delta for binary variables and *ε*^2^ for multi-group comparisons (Figure 1 D). Sex showed the strongest overall effect, particularly for SDNN (*ε*^2^ ≈ 0.40), followed by VLF (*ε*^2^ ≈ 0.22) and LF (*ε*^2^ ≈0.14), indicating substantial differences in cardiac autonomic regulation between men and women. Age group produced smaller but consistent effects, especially for LF and HF, whereas BMI and blood pressure categories showed minimal effect sizes across most HRV variables, with *ε*^2^ values typically below 0.05. These results indicate that, although conventional physiological descriptors influence HRV, their individual contribution is limited and insufficient to explain the full range of variability observed in the cohort.

When combinations of physiological descriptors were considered, effect sizes increased but remained moderate (Figure 1 E–F). Double-category comparisons revealed that interactions involving age contributed more to HRV discrimination than BMI or blood pressure, particularly for SDNN, LF, and HF, where *ε*^2^ values reached approximately 0.10–0.11 in some combinations. Triple-category comparisons produced slightly larger effects in selected cases, but no combination yielded large effect sizes across all HRV metrics. This pattern indicates that autonomic regulation in this population reflects a multidimensional structure in which several factors contribute simultaneously, rather than a simple hierarchy dominated by a single physiological variable.

A global ranking of HRV variables based on the number of significant comparisons and maximum effect size confirmed that LF, SDNN, and VLF were the most discriminative metrics across the evaluated factors (Figure 1 G). LF showed the highest number of significant results, followed by SDNN and VLF, whereas RMSSD and mean HR showed the lowest discriminative power. This finding suggests that long-term and spectral components of HRV capture broader aspects of autonomic regulation, while short-term measures may reflect more localized or context-dependent processes, resulting in lower discriminative capacity at the population level.

To further illustrate the interaction between demographic factors, HRV distributions were examined across sex and age ranges (Figure 1 H–I). LF values tended to decrease with increasing age, particularly in men, while women showed a broader dispersion across age groups. SDNN exhibited a similar pattern, with larger variability in younger groups and reduced dispersion in older individuals, although considerable overlap remained between categories. The persistence of overlap across all comparisons indicates that conventional descriptors such as age, sex, BMI, or blood pressure define only coarse groupings and fail to capture the full physiological heterogeneity present in the cohort.

The statistical characterization demonstrates that autonomic regulation in this population is highly heterogeneous and cannot be fully explained by single clinical or demographic variables. The moderate effect sizes observed across individual and combined descriptors indicate the presence of latent physiological substructures that are not directly observable through conventional categorizations. This multidimensional variability provides a strong rationale for the use of unsupervised approaches to identify coherent autonomic profiles, as explored in the following sections.

### 2.3 Exploratory clustering benchmark and selection of candidate partitions

To identify latent structure in the autonomic and physiological dataset without imposing predefined labels, an extensive exploratory clustering benchmark was performed across a large number of algorithmic configurations, feature representations, and parameter combinations. A total of 24,168 clustering experiments were executed, covering 12 clustering algorithms grouped into 8 methodological families, 38 different input spaces derived from alternative preprocessing and dimensionality-reduction strategies, and a wide range of cluster numbers. Only valid solutions passing consistency and quality filters were retained for subsequent analysis.

Figure 2 summarizes the global behavior of the exploration. The number of valid experiments obtained for each clustering family is shown in panel A, indicating that density-based, hierarchical, and model-based approaches generated the largest number of valid partitions, whereas community-graph and message-passing methods produced fewer acceptable solutions. This imbalance reflects both the intrinsic stability of certain algorithmic families and the stricter validity constraints imposed by graph-based and message-passing approaches.

**Figure 2:**
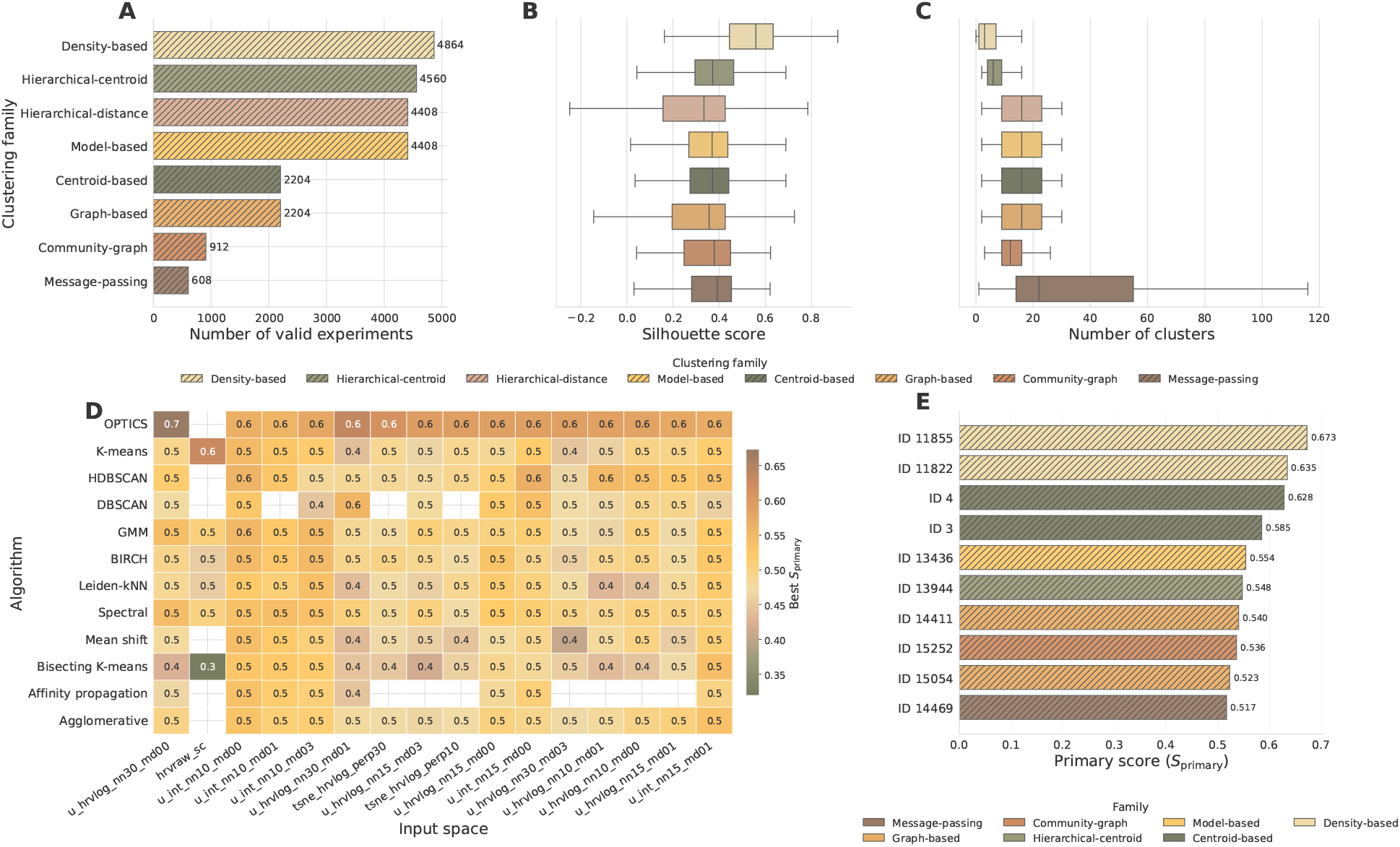
Exploratory benchmark of clustering strategies across multiple algorithms, feature representations, and parameter configurations. (A) Number of valid clustering experiments obtained for each clustering family after quality filtering. (B) Distribution of silhouette scores across families. (C) Distribution of the number of clusters generated by each family. (D) Heatmap showing the best primary score (*S*_primary_) obtained for each algorithm–input space combination, where input spaces correspond to alternative feature representations derived from HRV variables and integrated physiological descriptors. (E) Top 10 clustering strategies ranked by *S*_primary_, colored by clustering family. *S*_primary_ represents a composite internal validation score combining separation quality, cluster balance, and partition consistency.

The distribution of silhouette scores across families (Figure 2B) revealed clear differences in clustering performance. Density-based methods showed the highest median silhouette values, together with a relatively compact distribution, indicating robust separation in several feature spaces. Hierarchical and centroid-based algorithms exhibited moderate but stable performance, whereas graph-based and message-passing approaches displayed higher variability, suggesting stronger dependence on the chosen representation. These results indicate that no single algorithmic family consistently dominates across all input spaces, supporting the need for a large-scale exploratory strategy.

The number of clusters generated by each family (Figure 2C) further highlights methodological differences. Density-based methods tended to produce fewer clusters, often identifying compact structures together with noise points, whereas message-passing and graph-based approaches occasionally generated larger partitions. Hierarchical and centroid-based methods produced intermediate cluster numbers with relatively narrow distributions, indicating more predictable behavior across parameter settings. This diversity in cluster cardinality confirms that fixing the number of clusters a priori would have restricted the exploration and potentially hidden meaningful physiological subgroups.

To evaluate the interaction between algorithms and feature representations, the best primary score obtained for each algorithm–space combination is shown in Figure 2D. High scores were not restricted to a single algorithm, but instead appeared across multiple families depending on the input space. In particular, representations derived from UMAP projections of HRV variables and integrated physiological descriptors frequently yielded higher scores, indicating that dimensionality reduction played a critical role in revealing separable structure in the data. Density-based, centroid-based, model-based, and graph-based algorithms all produced competitive solutions under specific representations, reinforcing the importance of evaluating multiple algorithm–space combinations rather than relying on a single method.

The ten highest-scoring clustering strategies identified during the exploration are shown in Figure 2E and summarized in Table 1. These solutions span several algorithmic families, including density-based (OPTICS), centroid-based (K-means), model-based (GMM), hierarchical-centroid (Birch), graph-based (Spectral), community-graph (Leiden-KNN), and message-passing (Affinity propagation), indicating that highquality partitions can arise from fundamentally different clustering principles. Notably, the top-ranked solutions were consistently obtained in reduced or integrated feature spaces, particularly UMAP-based representations combining HRV-derived metrics and additional physiological descriptors. This result suggests that the structure of the data becomes more separable after nonlinear projection, likely because correlated physiological variables are compacted into lower-dimensional manifolds.

**Table 1:**
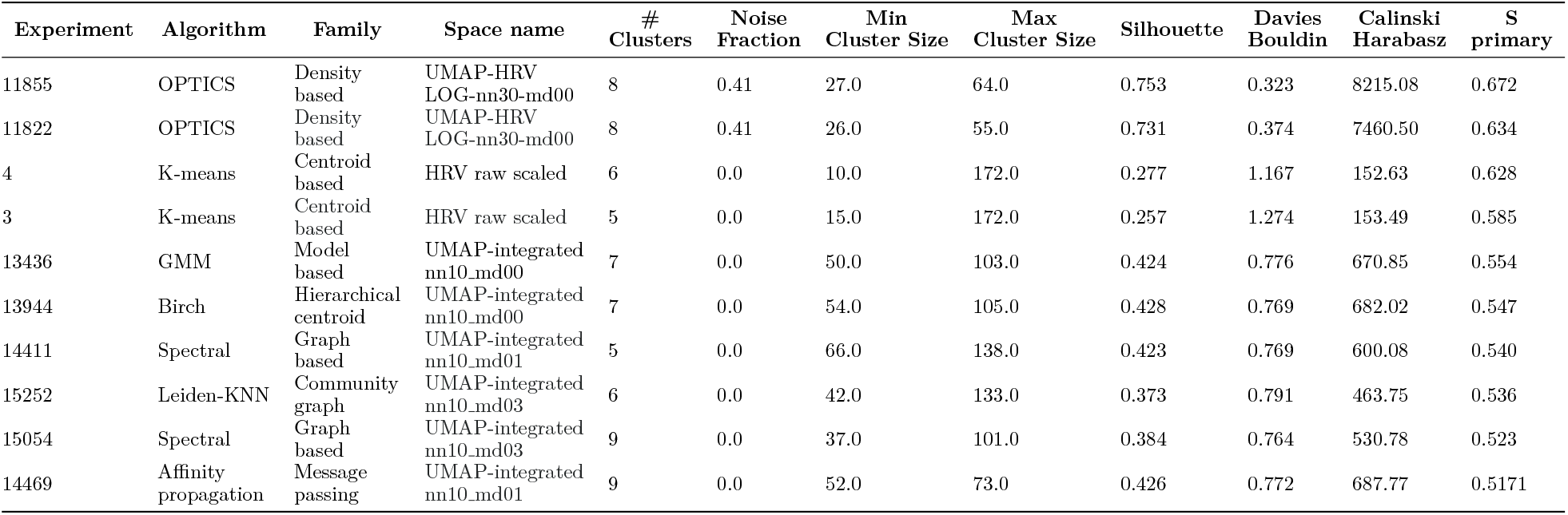
Summary of the top 10 best-performing clustering strategies identified during the exploration phase for candidate partition generation.

Although several strategies achieved similar primary scores, the selection of candidate partitions for detailed characterization required additional criteria beyond internal validation metrics. Table 1 shows that the highest-scoring solution (Experiment 11855, OPTICS) involved a relatively high noise fraction and strong dependence on density parameters, whereas centroid-based solutions produced very unbalanced cluster sizes despite good scores. Model-based and hierarchical-centroid approaches generated more stable partitions, but in some cases showed reduced separation compared to the best-performing density-based solutions. Graph-based and community-based methods, on the other hand, provided partitions with intermediate scores but more homogeneous cluster sizes and lower noise fractions.

Considering score, cluster balance, stability across representations, and interpretability of the resulting groups, experiment 15252 (Leiden-KNN, community-graph family, UMAP-integrated nn10 md03 space) was selected as the reference partition for detailed analysis. Although not the highest-scoring configuration, this solution provided a favorable compromise between separation quality, cluster size distribution, and robustness, producing six clusters without noise points and with balanced group sizes. In addition, the use of a community-detection algorithm in a reduced integrated space yielded partitions that were more consistent with the underlying physiological structure of the dataset, facilitating subsequent biological interpretation.

The selected partition was therefore used for the detailed cluster characterization described in the following section, where the autonomic, anthropometric, and cardiovascular profiles associated with each group were analyzed in depth.

Because the highest internal score does not necessarily correspond to the most useful physiological partition, final selection was based on a multi-criteria framework that considered internal separation metrics together with noise fraction, cluster-size balance, consistency across neighboring representations, and downstream interpretability in the original physiological variables. Under this framework, the six-cluster Leiden-KNN partition was retained as the reference solution because it offered the most favorable compromise between structural separation, balanced group sizes, absence of noise points, and physiological coherence. This partition was therefore used as the main descriptive solution for downstream characterization, while acknowledging that its robustness should be further examined in independent cohorts and dedicated stability analyses.

### 2.4 Physiological interpretation of the selected autonomic profiles

For clarity, the six clusters (C0–C5) are best interpreted as candidate autonomic profiles distributed along gradients of variability, cardiovascular load, and adiposity rather than as fixed disease categories (Figure 3A). This framing is important because the UMAP projection showed partial overlap and continuity between groups, suggesting that the detected phenotypes reflect structured physiological organization embedded within a broader continuum. Continuous variables contributing most strongly to cluster separation included SDNN, LF, VLF, and systolic blood pressure (Figure 3B), while blood pressure category, sex, and BMI category also differed across clusters (Figure 3C), supporting the physiological relevance of the partition..

**Figure 3:**
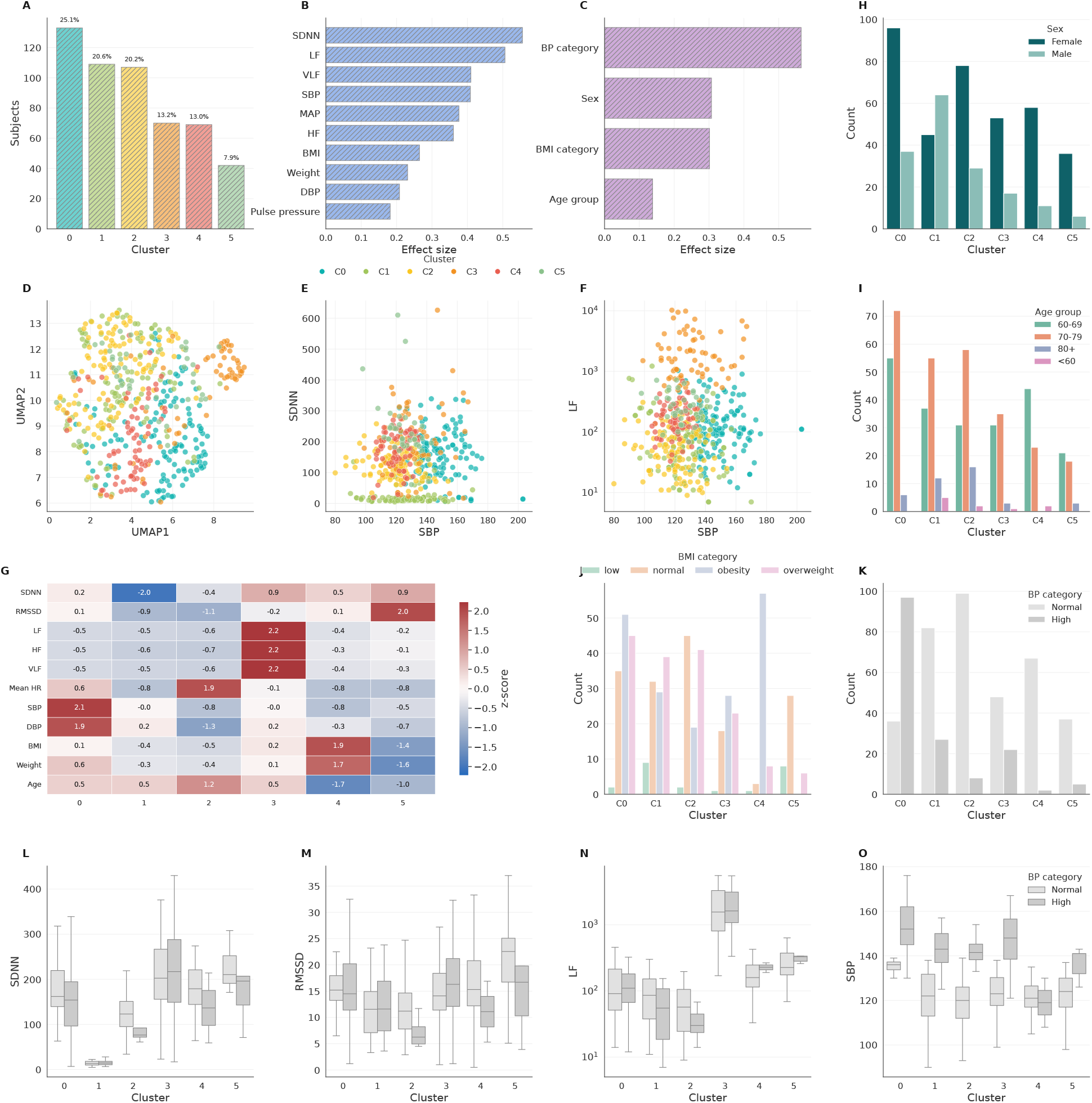
Physiological characterization of the selected clustering solution (strategy 15252). (A) Cluster sizes showing a balanced distribution of subjects across six clusters (C0–C5). (B) Ranking of continuous variables by effect size across clusters, highlighting HRV indices and blood pressure measures as the main contributors to separation. (C) Effect sizes for categorical variables, with strong differences for blood pressure category, sex, and BMI category. (D) UMAP projection colored by cluster, showing partially separated regions connected by gradual transitions. (E–F) Relationships between systolic blood pressure and SDNN or LF across clusters. (G) Standardized median profiles (z-scores) of physiological variables, showing distinct autonomic, cardiovascular, and anthropometric signatures. (H–K) Distribution of sex, age group, BMI category, and blood pressure category across clusters. (L–O) Boxplots of SDNN, RMSSD, LF, and SBP stratified by blood pressure category, showing that cluster differences persist within BP strata. Overall, the selected partition captures physiologically interpretable patterns of autonomic regulation in the cohort.

In the low-dimensional projection, subjects formed partially separated regions connected by gradual transitions (Figure 3D), suggesting that autonomic regulation in the cohort is organized along continuous gradients rather than discrete categories. This pattern is consistent with the concept that aging-related changes in autonomic control occur progressively and involve multiple interacting physiological dimensions. The relationship between systolic blood pressure and HRV indices further illustrates these gradients. Clusters differed not only in blood pressure levels but also in overall variability and spectral components, as shown by the distributions of SDNN and LF (Figures 3E–F). Groups with higher blood pressure tended to exhibit reduced variability or altered spectral balance, a pattern compatible with less favorable autonomic regulation and greater cardiovascular load.

The standardized median profiles highlight the distinct physiological signatures of each cluster (Figure 3G). One group showed markedly reduced SDNN and RMSSD values, indicating limited variability and suggesting reduced autonomic flexibility. Another cluster displayed elevated LF, HF, and VLF components together with high SDNN, consistent with a profile of high overall variability and strong autonomic modulation. A different group combined higher BMI and body weight with moderate HRV values, while another showed elevated blood pressure with relatively preserved variability. These combinations indicate that the clusters represent coherent physiological patterns involving autonomic, cardiovascular, and anthropometric variables rather than differences driven by a single parameter.

The distribution of categorical variables reinforces this interpretation. Sex composition varied across clusters (Figure 3H), and age-group distribution showed that older individuals were more frequent in clusters characterized by lower variability and higher blood pressure (Figure 3I). Body mass index categories were unevenly distributed across clusters (Figure 3J), with higher BMI values being more common in groups showing altered HRV profiles, consistent with the known association between metabolic status and autonomic regulation. Blood pressure categories also differed markedly between clusters (Figure 3K), confirming that the partition captures clinically relevant differences in cardiovascular condition.

Importantly, differences between clusters remained evident even after stratifying by blood pressure category. Comparisons of SDNN, RMSSD, and LF showed clear separation between clusters within the same BP category (Figures 3L–N), indicating that the clustering is not simply reflecting blood pressure classification but captures additional variability in autonomic regulation. Similarly, systolic blood pressure itself differed between clusters within the same BP group (Figure 3O), supporting the idea that autonomic control, cardiovascular load, and metabolic status represent partially independent but interacting dimensions.

Taken together, these results suggest that aging-related autonomic heterogeneity is not random, but organized into recurrent multivariable configurations combining HRV amplitude, hemodynamic load, body composition, and demographic structure. Accordingly, the selected clustering strategy should be interpreted as a data-driven descriptive representation of latent autonomic organization within this cohort, rather than as a definitive taxonomy of validated physiological phenotypes.

## 3 Conclusions

In conclusion, HRV in older adults from the Magallanes region is not adequately explained by isolated clinical descriptors alone, but instead reflects a structured multidimensional physiological organization. Integrated unsupervised analysis identified six candidate autonomic profiles that differed in variability, hemodynamic burden, body-composition patterns, and demographic structure. These findings support the use of multi-variable approaches for interpreting autonomic heterogeneity in aging populations. However, the detected profiles should be regarded as data-driven hypotheses of latent physiological organization that now require longitudinal assessment, formal stability testing, and external validation before clinical or translational use.

The descriptive and correlational analyses revealed that HRV variables, particularly spectral and long-term variability components such as LF, VLF, and SDNN, captured broader physiological variation than shortterm metrics alone. At the same time, the relatively weak and partial associations observed between HRV, anthropometric variables, and blood pressure measurements suggest that autonomic function in older adults is shaped by multiple interacting domains rather than by a single dominant physiological factor. This multidimensional structure supports the need for integrative analytical strategies when interpreting HRV in aging populations.

The main finding of this study is that autonomic regulation in older adults from southern Chile is better represented as an integrated multivariable organization than as the additive effect of isolated descriptors such as age, sex, BMI, or blood pressure. Although these conventional factors explained part of the observed variability, they did not adequately capture the latent physiological structure revealed by the unsupervised analysis.

From a physiological perspective, the identified profiles suggest that autonomic regulation in older adults is better understood as a continuum of partially overlapping physiological states, ranging from relatively preserved variability to profiles compatible with reduced autonomic flexibility and greater cardiovascular burden.

This study has several strengths, including the relatively large cohort size, the integration of multiple physiological domains, the use of a transparent preprocessing workflow, and the systematic exploration of clustering strategies across diverse feature spaces and algorithmic families. However, several limitations should temper interpretation. The cross-sectional design precludes causal or prognostic inference, the convenience-sampling strategy limits representativeness, and the detected profiles were derived from internal structure within a single cohort. In addition, because short-term spectral HRV indices are not physiologically univocal, cluster interpretation should rely on multivariable patterns rather than on any single component in isolation. Accordingly, the present findings should be viewed as descriptive and hypothesis-generating until supported by formal stability analyses and external validation.

In summary, this study establishes a detailed descriptive framework for interpreting HRV in older adults from an extreme southern region and demonstrates that autonomic variability in this population is multidimensional, structured, and only partially captured by traditional clinical descriptors. These findings provide an empirical basis for future longitudinal, inferential, and predictive studies aimed at improving the characterization of autonomic health in aging populations.

## 4 Methods and implementation strategies

A multi-stage analytical pipeline was implemented to characterize autonomic regulation in older adults using heart rate variability (HRV) together with demographic, anthropometric, and cardiovascular descriptors. The overall workflow of the study is summarized in Figure 4. The proposed strategy integrates standardized physiological acquisition, reproducible data preprocessing, statistical characterization of the cohort, and an extensive unsupervised clustering benchmark aimed at identifying latent patterns of autonomic regulation without predefined labels.

**Figure 4:**
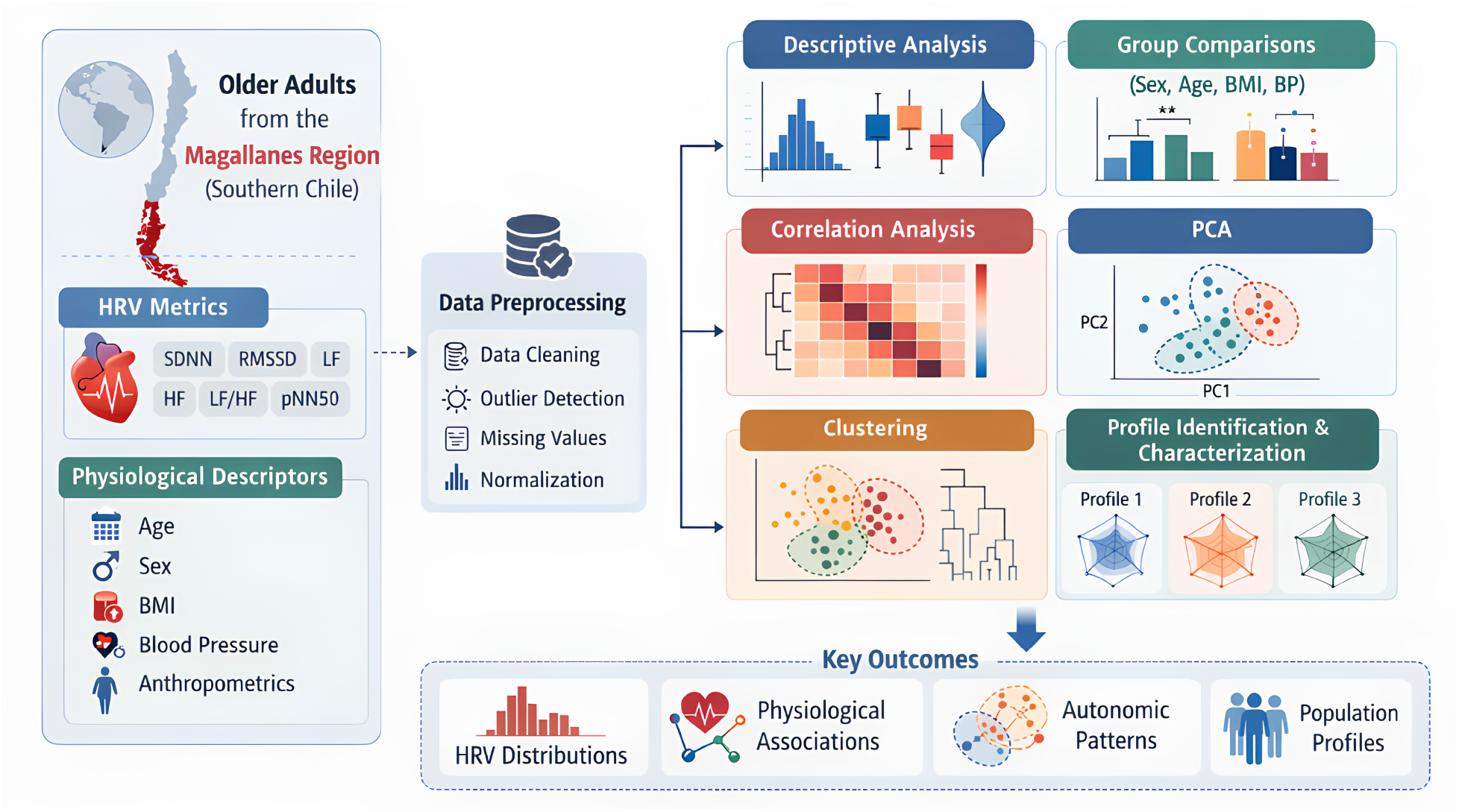
Overview of the analytical pipeline used for the statistical characterization of autonomic regulation in older adults from the Magallanes region (southern Chile). The workflow integrates heart rate variability (HRV) metrics together with demographic, anthropometric, and cardiovascular descriptors. After standardized data acquisition, a preprocessing stage including data cleaning, outlier detection, missing value handling, and normalization was applied to construct a consistent dataset. The processed data were then explored using descriptive statistics, group comparisons, correlation analysis, and dimensionality reduction. An extensive unsupervised clustering benchmark was performed across multiple feature spaces and algorithms to identify latent physiological structure. Selected clustering solutions were subsequently used for profile identification and characterization, allowing the detection of distinct autonomic regulation patterns within the studied population.

First, HRV metrics and physiological descriptors were obtained under controlled experimental conditions in a community-dwelling population of older adults from the Magallanes region in southern Chile. Raw data were then processed using a reproducible preprocessing pipeline including data cleaning, normalization, consistency checks, and generation of quality control indicators to ensure numerical reliability before downstream analyses.

After preprocessing, the dataset was characterized through descriptive statistics, group comparisons, and correlation analyses in order to evaluate the distribution of autonomic and physiological variables and to identify relevant associations between HRV metrics and clinical descriptors. Dimensionality reduction techniques were additionally explored to visualize the structure of the data in reduced feature spaces and to assess the presence of potential latent organization.

To investigate whether the cohort could be partitioned into physiologically meaningful subgroups, an extensive unsupervised clustering benchmark was performed across multiple feature representations, dimensionality reductions, and clustering algorithms. Candidate clustering solutions were evaluated using internal validation metrics and partition descriptors, allowing comparison of alternative representations of autonomic regulation without imposing predefined categories.

Finally, selected clustering solutions were used to define participant profiles, which were subsequently characterized according to HRV metrics, demographic variables, and physiological descriptors. This procedure allowed the identification of distinct autonomic regulation patterns within the studied population and provided a data-driven characterization of autonomic variability in older adults.

### 4.1 Study design and participants

An observational cross-sectional study design was used to investigate autonomic function in community-dwelling older adults aged 60 years and older from the Magallanes and Chilean Antarctic region in southern Chile. All measurements were obtained at a single assessment time point, allowing the evaluation of associations between demographic, anthropometric, cardiovascular, and autonomic variables without longitudinal intervention.

Participants were recruited through non-probabilistic convenience sampling from the local community. A total of 530 subjects were included in the study. Eligibility criteria required participants to be 60 years of age or older, permanently residing in the Magallanes region, and presenting a Karnofsky Performance Status score of at least 60%, ensuring sufficient functional autonomy to complete the study procedures. Individuals were excluded if they had a diagnosis of diabetic neuropathy, pacemaker implantation, clinical depression, dementia, or severe cognitive or motor impairment.

Additional exclusion criteria included the use of beta-blockers at the time of evaluation, consumption of stimulant substances or medication within 12 hours prior to cardiac assessment, or any motor limitation preventing correct execution of the measurements.

All participants received detailed information about the study objectives, procedures, risks, and responsibilities prior to enrollment, and all subjects provided written informed consent before participation.

The study protocol was approved by the Ethics Committee of the University of Chile (ACTA N°029-18/05/2022) and the Ethics Committee of the University of Magallanes (N°008/SH/2022), and all procedures were conducted in accordance with the Declaration of Helsinki for research involving human subjects.

### 4.2 Experimental protocol and data acquisition

All evaluations were performed at the Centro Asistencial Docente y de Investigación (CADI-UMAG), a clinical and research facility affiliated with the University of Magallanes in Punta Arenas, Chile. Measurements were obtained in a controlled laboratory environment between 9:00 and 11:00 a.m. in order to minimize circadian influences on cardiovascular and autonomic variables.

Environmental conditions were standardized for all participants. Room temperature was maintained at approximately 20°C to ensure thermal comfort and to reduce the potential influence of thermoregulatory responses on autonomic function. Artificial white light was used to provide constant illumination, and all assessments were conducted in a quiet and private setting to minimize external interference.

Body composition was assessed using multi-frequency bioelectrical impedance analysis with a Tanita BC-558 Ironman Segmental Body Composition Monitor (Tanita Ironman, Arlington Heights, IL, USA). Measurements included body weight, height, body mass index (BMI), body fat percentage, lean mass, total body water, and bone mass. Participants were instructed to fast for at least 4 hours before the evaluation, avoid strenuous exercise and alcohol consumption for at least 12 hours, and empty their bladder prior to the measurement. All measurements were performed with participants barefoot and wearing light clothing.

Resting systolic and diastolic blood pressure were measured using an automated Omron® monitor to characterize baseline cardiovascular status. To ensure safe and reliable autonomic assessment, only participants presenting blood pressure values below 140/90 mmHg at the time of evaluation were included in the heart rate variability recording procedure.

Cardiac autonomic activity was assessed from continuous R–R interval recordings obtained using the Polar Team2 system (Polar®, Finland). Participants remained seated at rest during the recording procedure. After a stabilization period, R–R intervals were continuously monitored, and the final portion of the resting period was used for analysis. A stable 5-minute segment was extracted for heart rate variability evaluation following previously validated procedures.

Participants were instructed to maintain spontaneous breathing during the recording and to avoid unnecessary movement or speaking. All recordings were visually inspected to ensure signal quality before further processing.

### 4.3 Heart rate variability analysis

Heart rate variability (HRV) analysis was performed from R–R interval recordings using Kubios HRV® software (Kubios Oy, Kuopio, Finland), following established guidelines for short-term HRV assessment. Prior to analysis, all recordings were visually inspected and automatically corrected for artifacts and ectopic beats using the software’s built-in filtering algorithm. Recordings containing more than 3% of corrected beats were excluded from further analysis, in accordance with recommended standards for HRV studies (Malik, 1996).

Time-domain variables included mean R–R interval, mean heart rate, the standard deviation of normal R–R intervals (SDNN), and the root mean square of successive differences between adjacent R–R intervals (RMSSD). SDNN was considered an indicator of overall autonomic modulation, whereas RMSSD was used as a marker of parasympathetic activity (Buchheit et al., 2010).

Frequency-domain analysis was performed using fast Fourier transform as implemented in Kubios HRV®. The following spectral components were obtained: very-low-frequency power (VLF), low-frequency power (LF), and high-frequency power (HF). HF power was interpreted as an index of parasympathetic modulation, while LF power was considered to reflect baroreflex-related autonomic regulation. VLF power was included due to its reported association with long-term regulatory and stress-related processes.

To further characterize autonomic balance, composite indices were calculated using Kubios HRV®. The Parasympathetic Nervous System Index (PNS index) was derived from mean R–R interval, RMSSD, and the Poincaré plot SD1 index, all expressed in normalized units. The Sympathetic Nervous System Index (SNS index) was derived from mean R–R interval, the Baevsky Stress Index, and the Poincaré plot SD2 index. In addition, the Stress Index (SI), based on Baevsky’s formulation, was included as an indicator of autonomic load and sympathetic predominance (Berntson et al., 1997; Rajendra Acharya et al., 2006; Baevsky and Chernikova, 2017).

All HRV variables used in subsequent analyses were extracted from the same 5-minute stable segment obtained under resting conditions.

### 4.4 Dataset construction and preprocessing

All preprocessing steps were performed using custom scripts written in Python (version X.X), using the libraries `pandas, numpy`, and `scipy`. The raw dataset was obtained from the original spreadsheet file and processed through a standardized pipeline designed to ensure numerical consistency, detect potential recording errors, and generate quality control indicators before statistical and clustering analyses.

Raw data were first imported from the original Excel file and converted into a tabular format. Numeric variables were cleaned to correct formatting inconsistencies commonly found in manually recorded datasets, including removal of extra spaces, correction of decimal separators, and conversion of malformed numeric strings into valid floating-point values. Invalid or empty values were converted to missing values.

After numeric normalization, derived variables were recomputed to verify internal consistency. Body mass index (BMI) was recalculated from height and weight, mean arterial pressure was recomputed from systolic and diastolic blood pressure, and heart rate was recomputed from the mean R–R interval. Differences between recorded and recomputed values were stored as diagnostic variables and later used as quality indicators.

Range validation was applied to physiological variables using predefined acceptable limits for anthropometric, cardiovascular, and HRV parameters. Values outside the expected ranges were not automatically removed but were flagged to allow later inspection and filtering if necessary.

Additional consistency checks were performed to detect potential errors in height–BMI relationships, heart rate–R–R interval correspondence, and other physiological constraints. For each subject, the number of detected inconsistencies was recorded, generating a global quality control score.

All preprocessing steps were logged automatically, including the initial dataset dimensions, transformations applied, and final dataset size. Intermediate diagnostic tables containing detected inconsistencies and range violations were exported for traceability.

The final cleaned dataset was exported in both CSV and Excel formats and used as the input for all subsequent statistical analyses and unsupervised learning procedures.

### 4.5 Statistical analysis

All statistical analyses were performed using Python (version X.X) with the libraries `pandas, numpy, scipy`, and `statsmodels`. The objective of this stage was to characterize the physiological dataset, evaluate differences between demographic groups, and identify patterns that could guide the subsequent unsupervised clustering analysis.

Descriptive statistics were computed for all variables, including mean, standard deviation, median, interquartile range, minimum, and maximum values. These analyses were performed for the complete cohort and stratified by sex and age categories in order to identify potential differences in autonomic and cardiovascular variables.

Normality of continuous variables was evaluated using the Shapiro–Wilk test. Because several variables did not follow a normal distribution, both parametric and non-parametric approaches were considered depending on the characteristics of each variable.

Comparisons between groups were performed using Student’s t-test or Mann–Whitney U test for two-group comparisons, and one-way ANOVA or Kruskal–Wallis test for multiple-group comparisons, as appropriate. When significant differences were detected, post-hoc analyses with multiple comparison correction were applied.

To facilitate interpretation of physiological patterns, continuous variables such as age, body mass index, and blood pressure were additionally categorized into clinically meaningful ranges. Group-based comparisons were then repeated using these categories in order to evaluate how autonomic variables varied across physiological conditions.

Correlation analyses were performed using Pearson or Spearman correlation coefficients depending on the distribution of the variables. Correlation matrices were used to evaluate relationships between anthropometric, cardiovascular, and HRV variables and to identify redundant or strongly associated features.

In addition, multivariable exploratory analyses were conducted to examine the combined behavior of physiological variables across the cohort. These analyses were not intended for predictive modeling but rather to support the identification of relevant variable groups to be used in the unsupervised clustering stage.

All statistical results, including summary tables, group comparisons, and correlation matrices, were automatically exported to ensure reproducibility and traceability of the analysis.

### 4.6 Unsupervised clustering strategy

To explore latent physiological structure in the cohort without imposing predefined labels, an unsupervised clustering benchmark was implemented as a systematic exploratory procedure. At this stage, the objective was not to define the final participant profiles, but rather to evaluate a broad range of clustering configurations across multiple feature representations and algorithmic families in order to identify potentially meaningful partitions for subsequent analysis.

Clustering was performed on several complementary input spaces derived from the preprocessed dataset. These included: i) raw HRV variables after imputation and standardization, ii) log-transformed HRV variables after imputation and standardization, iii) standardized physiological and anthropometric variables, and iv) an integrated space combining log-transformed HRV variables with continuous physiological descriptors. To further evaluate whether lower-dimensional representations improved cluster structure, additional embedding spaces were generated from selected datasets using principal component analysis (PCA), t-distributed stochastic neighbor embedding (t-SNE), and uniform manifold approximation and projection (UMAP), when available. In this way, clustering was explored both in the original standardized feature spaces and in alternative reduced-dimensional manifolds.

A broad panel of clustering algorithms was evaluated in order to cover different assumptions about data structure. The exploratory benchmark included centroid-based methods (*k*-means and bisecting *k*-means), model-based clustering (Gaussian mixture models), hierarchical and distance-based methods (agglomerative clustering and BIRCH), density-based methods (DBSCAN, HDBSCAN, OPTICS, and MeanShift), graph-based methods (spectral clustering), message-passing methods (affinity propagation), and community detection on *k*-nearest neighbor graphs using the Leiden algorithm. This design was intended to avoid privileging a single notion of cluster structure, given the heterogeneity expected in physiological and autonomic data.

For partition-based methods requiring a predefined number of clusters, the number of clusters was systematically varied from 2 to 30. For Gaussian mixture models, multiple covariance structures were evaluated. For agglomerative clustering, several linkage criteria were considered. Density-based methods were explored across a grid of neighborhood size, minimum cluster size, and density sensitivity parameters. For MeanShift, bandwidth was estimated automatically under different quantile settings. For Leiden community detection, clustering resolution and graph neighborhood size were varied systematically. This strategy generated a large set of candidate clustering solutions spanning multiple representations and modeling assumptions.

Before clustering, all numerical input spaces were median-imputed and standardized to zero mean and unit variance. Log-transformation was specifically applied to skewed HRV variables to reduce the influence of extreme values and improve comparability across autonomic metrics with different numerical scales. Dimensionality reduction methods were applied only after imputation and scaling, and the resulting embeddings were treated as alternative exploratory input spaces rather than as definitive representations.

Each clustering experiment was executed independently and stored with its corresponding metadata, including algorithm, algorithm family, input space, parameter configuration, and cluster labels. In addition, each solution was summarized by basic partition descriptors such as the number of detected clusters, cluster size distribution, and fraction of observations assigned to noise when applicable. All label assignments were exported as separate files to support downstream inspection, profile characterization, and validation analyses.

Because the present subsection focuses exclusively on exploratory benchmarking, cluster quality was assessed only through internal unsupervised criteria. For each valid clustering solution, silhouette score, Calinski– Harabasz index, and Davies–Bouldin index were computed, together with the fraction of noise points for algorithms that allowed unassigned observations. These metrics were used to generate preliminary rankings of candidate solutions and to compare performance across algorithms, algorithmic families, and input spaces. However, the final selection of relevant clustering solutions, together with the physiological characterization and validation of the identified participant profiles, was reserved for a subsequent analysis stage.

### 4.7 Exploratory clustering design and multi-criteria selection of physiologically meaningful partitions

The identification of latent patterns of autonomic regulation was approached as an exploratory unsupervised learning problem in which no single feature representation, dimensionality reduction, or clustering algorithm could be assumed to be optimal a priori. The objective of the study was not to obtain a geometrically optimal partition of the data, but to detect physiologically meaningful subgroups characterized by coherent heart rate variability profiles and interpretable relationships with demographic and cardiovascular descriptors. Because different representations of the data may reveal different aspects of autonomic regulation, a large-scale clustering exploration was performed across multiple feature spaces, embedding strategies, and clustering algorithms, followed by a multi-criteria selection procedure designed to prioritize partitions that satisfied structural validity, physiological coherence, and descriptive interpretability simultaneously.

The exploratory stage generated a large set of candidate partitions obtained from heterogeneous representations of the data. These representations included spaces derived from heart rate variability variables, spaces defined by physiological and demographic descriptors, and integrated representations combining both domains. Dimensionality-reduced embeddings were constructed using linear and non-linear methods in order to evaluate the effect of geometric transformations on cluster structure. Each representation was analyzed using clustering algorithms belonging to different methodological families, including centroid-based, hierarchical, density-based, graph-based, community detection, and probabilistic approaches. This strategy produced a diverse collection of candidate partitions obtained under distinct geometric assumptions without imposing a predefined model of the latent structure.

Let 𝒫 = {*P*_1_, *P*_2_, …, *P*_*M*_} denote the set of all candidate partitions generated during the exploration. Each partition *P*_*i*_ is characterized by the feature space used to construct the representation, the clustering algorithm, the algorithm family, the number of clusters *k*_*i*_, the fraction of observations labeled as noise *η*_*i*_, internal validation metrics, and the assignment of subjects to clusters. Because internal clustering metrics evaluate geometric separation but do not guarantee physiological plausibility, candidate partitions were evaluated using a multi-criteria framework combining structural quality, intra-cluster physiological coherence, preservation of physiological relationships, and interpretability with respect to demographic and cardiovascular descriptors.

Candidate partitions were first subjected to filtering constraints in order to remove trivial, degenerate, or excessively fragmented solutions. Only partitions with a moderate number of clusters were retained. The number of clusters was restricted to the interval

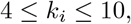

with preference for partitions satisfying

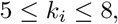

which provided sufficient granularity to capture heterogeneous autonomic profiles while preserving adequate sample size within each group. Partitions producing extremely small clusters were excluded by requiring that the smallest cluster contained at least ten subjects. Solutions with excessive noise were removed by imposing an upper bound on the fraction of observations labeled as noise,

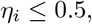

and partitions with negligible structural separation were discarded by requiring non-negative Silhouette values. Degenerate solutions dominated by a single cluster were avoided by constraining the maximum cluster proportion,

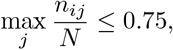

where *n*_*ij*_ denotes the number of subjects in cluster *j* of partition *i*, and *N* the total sample size.

For the partitions that satisfied these constraints, a structural score was computed from internal validation metrics. Let 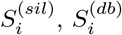 and 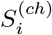 denote normalized versions of the Silhouette, inverted Davies–Bouldin, and Calinski–Harabasz indices. A penalty term was introduced to down-weight solutions outside the preferred cluster range. The structural score was defined as

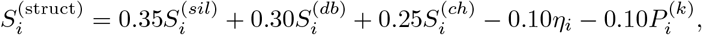

where the cluster penalty was defined as

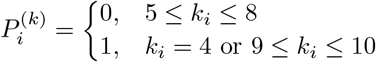

and partitions outside the interval 4 ≤ *k*_*i*_ ≤ 10 were removed during the filtering stage.

To evaluate whether a partition produced physiologically coherent groups, the dispersion of heart rate variability variables was analyzed in the original HRV domain, independently of the feature space used to generate the partition. For each HRV variable *v*, the reduction of variance induced by partition *P*_*i*_ was computed as

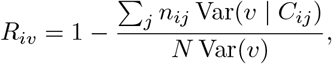

where *C*_*ij*_ denotes cluster *j* of partition *i*. In addition, cluster compactness was evaluated using the interquartile range. The intra-cluster physiological coherence score was defined as

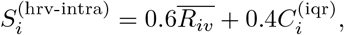

with

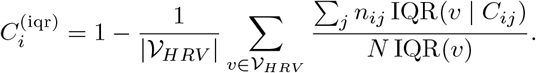

Physiologically meaningful partitions should also preserve coherent relationships between HRV variables reflecting coordinated autonomic regulation. Instead of enforcing equality with the global correlation matrix, relational consistency was evaluated by measuring the proportion of HRV variable pairs that preserved the direction of association observed in the reference population. Let ℰ denote the set of HRV variable pairs considered, and let *r*_*i,uv*_ denote the correlation within partition *P*_*i*_. The relational score was defined as

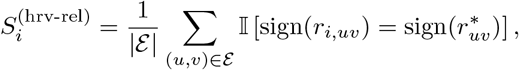

where 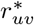 denotes the reference direction of association computed in the full cohort.

Interpretability with respect to demographic and cardiovascular descriptors was evaluated by measuring the magnitude of differences between clusters for external variables including age, sex, anthropometric measures, and blood pressure indicators. For each external variable *x*, an effect size Δ_*ix*_ was computed using an appropriate statistic depending on the variable type. The descriptive score was defined as

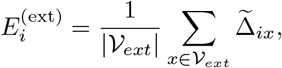

where 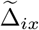 denotes the normalized effect size. To avoid trivial partitions driven by a single variable, a dominance penalty was defined as

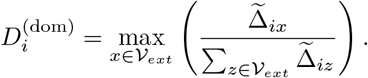

The descriptive interpretability score was defined as

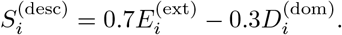

The primary ranking score combined structural validity and physiological criteria,

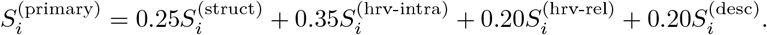

Because the exploratory stage produced many highly similar partitions, the final selection enforced diversity between clustering algorithms, algorithm families, feature spaces, and the number of clusters. Partitions were ranked according to the primary score, and the final Top-*K* set was obtained by sequential selection while preventing dominance of any single algorithm, representation, or cluster configuration. This procedure ensured that the selected solutions represented distinct and physiologically plausible views of the latent structure of autonomic regulation.

## Informed Consent Statement

All participants received detailed information regarding the study objectives, procedures, and potential implications. Informed consent was obtained to ensure ethical compliance and participant autonomy.

## AI use statement

The authors used artificial intelligence-assisted tools exclusively for editorial support, including language polishing, grammar correction, and improvement of textual clarity. No artificial intelligence system was used to generate the study design, perform the statistical analyses, produce the reported results, or determine the scientific interpretations and conclusions. All outputs produced with editorial assistance were critically reviewed, edited, and validated by the authors, who assume full responsibility for the final content of the manuscript.

## Author contributions statement

Conceptualization, DM-O, MC-A, DM-C, MM-C, CN-E; Data curation, D M-O, MC-A; Investigation, DM-O, MC-A, DM-C, MM-C, CN-E; Methodology, DM-O, MC-A; Supervision, DM-O, CN-E; Formal analysis, DM-O; Visualization, DM-O, MC-A; Writing–original draft, DM-O, MC-A, DM-C, MM-C, CN-E; Writing–review & editing, DM-O, MC-A, DM-C, MM-C, CN-E. All authors have read and agreed to the published version of the manuscript.

## Acknowledgments

This work was funded by ANID Proyecto Fondecyt Regular Nº1250474 and by the Innovation Fund for Competitiveness of the Regional Government of Magallanes and Chilean Antarctica (BIP Code 40042452-0). DM-O acknowledges funding from FONDECYT Iniciación 11250295. DM-O gratefully acknowledges support from the Center for Biotechnology and Bioengineering - CeBiB (PIA project FB0001, Conicyt, Chile). CN-E was funded by the Chilean National Agency for Research and Development (ANID) under project Fondecyt Regular N°1250474. The authors sincerely thank the older adults who participated in this study for their time, commitment, and valuable contribution. The authors also acknowledge the Centro Asistencial Docente e Investigación of the University of Magallanes (CADI-UMAG) for kindly providing the facilities used for participant assessments and study procedures.

## Code and data availability

The code used for data preprocessing, statistical analysis, clustering exploration, and figure generation is publicly available in a GitHub repository: https://github.com/nim-ach/heart_variability_rate_profiles.

The processed dataset and all derived summary tables generated during the analyses are also accessible through the same repository, or via an associated public data repository linked therein.

Due to ethical and privacy considerations associated with human participant data, the raw dataset is not publicly available. However, it can be made available by the authors upon reasonable request and subject to appropriate data access agreements.

